# Altering dosage of meiotic crossover-associated RING finger proteins affects crossover number and interference in Drosophila

**DOI:** 10.64898/2026.02.18.706578

**Authors:** Emerson Frantz, Priscila Santa Rosa, Susan McMahan, Jeff Sekelsky

## Abstract

Crossovers play a critical role in ensuring correct reductional segregation of homologous chromosomes in the first meiotic division. Crossing over is initiated by formation of DNA double-strand breaks (DSBs), but the number of DSBs is greater than the number of crossovers. Which recombination sites become crossovers, versus being repaired as non-crossovers, is not random, but is subject to several crossover patterning phenomena, including crossover assurance and crossover interference. One current model for crossover designation proposes that crossover-associated RING finger proteins (CORs) undergo the biophysical process of coarsening, in which larger accumulations continue to get larger and smaller accumulations go away. Genetic and cytological studies of the three CORs in *Drosophila melanogaster*, Vilya, Narya, and Nenya, are consistent with this model. In females heterozygous for a deletion of *vilya*, fewer double-crossovers are observed. Conversely, crossovers are elevated in females carrying a duplication of *vilya* and in females coordinately overexpressing *Vilya*, Narya, and Nenya. These findings support a model in which crossover designation occurs through coarsening of COR proteins within the synaptonemal complex.

## Introduction

Crossovers between homologous chromosomes facilitate reductional segregation in meiosis. Meiotic recombination is initiated by the introduction of DNA doublestrand breaks (DSBs), but the number of DSBs made is greater than the number of crossovers. Crossover patterning processes (reviewed in Pazhayam et al. (2021)) determine which sites are repaired to give crossovers. Perhaps the most important patterning outcome is crossover assurance, which dictates that each pair of homologs usually has at least one crossover per meiosis, regardless of chromosome size (Owen, 1949). Crossover assurance is important because chiasmata resulting from crossovers provide physical linkages that lead do stable biorientation of homologous chromosomes on the meiotic spindle, thereby promoting reductional segregation. Consequently, reductions in crossing over are associated with high rates of meiotic chromosome nondisjunction.

Another aspect of crossover patterning is interference, originally described by Sturtevant (1913) as the tendency of the *X* chromosomes in *Drosophila* to have only one crossover per meiosis. Subsequent definitions of interference include a significant reduction in double-crossovers (DCOs) in adjacent chromosome intervals compared to the number expected if the two intervals are independent of one another, or larger spacing between crossovers than expected in cases where there are two crossovers in one meiosis. Although interference is widespread, occurring in plants, fungi and metazoa, the reasons for its existence have been less clear (reviewed in Berchowitz and Copenhaver (2010); Otto and Payseur (2019)).

Several models have been proposed to explain how crossover patterning is achieved. One recent model posits that assurance and interference result from the biophysical process of coarsening (Morgan et al., 2021; Zhang et al., 2025). In this model, crossover designation factors are distributed within the synaptonemal complex (SC), a protein structure that assembles between paired homologous chromosomes. Rog *et al*. (2017) presented observations and experiments that suggest the SC is phase-separated, and Zhang *et al*. (2018) proposed that crossover designation occurs by diffusion of proteins within the SC. In the coarsening model, crossover designation proteins are initially distributed through the SC but are enriched at sites where DSBs are made to initiate recombination. Surface tension forces drive the accumulation of crossover designation factors at one or a few DSBs sites per SC, with other sites becoming depleted for these factors. This model explains crossover assurance, provided an SC has sufficient crossover designation factor to make at least one accumulation. If there is more of the designation factor(s), then additional designations can occur along a a single SC. These would likely be widely separated, resulting in the phenomenon of crossover interference.

In some species, crossover-associated RING finger proteins (CORs) exhibit behaviors consistent with them being crossover designation factors that accumulate through coarsening. These include HEI10 in *Arabidopsis thaliana* (Morgan et al., 2021) and ZHP-3 and ZHP-4 in *Caenorhabditis elegans* (Zhang et al., 2025). The *Drosophila melanogaster* genome encodes three CORs, named Vilya, Narya, and Nenya (Lake et al., 2015). Vilya physically interacts with Mei-P22, the noncatalytic subunit of the enzyme that generates meiotic DSBs, and is itself required for DSB formation. Studies of HA-tagged Vilya found that it is initially localized throughout the central region of the SC (Lake et al., 2015). As meiosis progresses, Vilya becomes less intense along the SC and more intense at foci that are similar in number to the average number of crossovers per meiosis. The requirement for Vilya in making DSBs precluded direct genetic tests of whether it has a later function in crossover designation or execution.

A study of Narya and Nenya by Lake *et. al* (2019) concluded that the *narya* gene arose from a duplication of *nenya* in the ancestor to the melanogaster species subgroup, and that loss of both Narya and Nenya results in failure to make meiotic DSBs. An in-frame deletion allele of *narya* that is predicted to delete five residues, including the last cysteine of the RING finger domain, is competent to make DSBs, but not to make crossovers when *nenya* is simultaneously knocked down. Together, these data show that DSB formation requires Vilya plus either Narya or Nenya. Studies of a separation-of-function allele of *narya* suggest that Narya or Nenya is also required to make crossovers, and that function requires an intact RING domain, though it is unknown whether ubiquitin ligase activity is required.

The involvement of Drosophila CORs in DSB formation makes it difficult to assess functions in crossover designation. We therefore explored the relationship between COR dosage and crossover number by assaying crossover numbers and crossover interference in flies heterozygous for a deletion of *vilya* and in flies overexpressing Vilya, Narya, and Nenya. Reducing Vilya dosage resulted in a significant decrease in the occurrence of double-crossover chromosomes. Conversely, whereas coordinate over-expression of all three CORs led to a significant increase in crossovers. These results, together with previously published data, support a role for Vilya, Narya, and Nenya in meiotic crossover designation through coarsening.

## Results and Discussion

### Reducing Vilya dosage leads to increased interference

To address whether reduced COR dosage impacts meiotic crossing over in Drosophila, we first assayed crossovers in females heterozygous for a deletion that removes *vilya*. In wild-type females, the genetic length of the interval assayed was 28.4 cM (Fig. 1A). In females heterozygous for a *vilya* deletion, the number of crossovers was not significantly changed (27.2 cM; 7666 progeny; *p* = 0.86).

**Figure 1.**
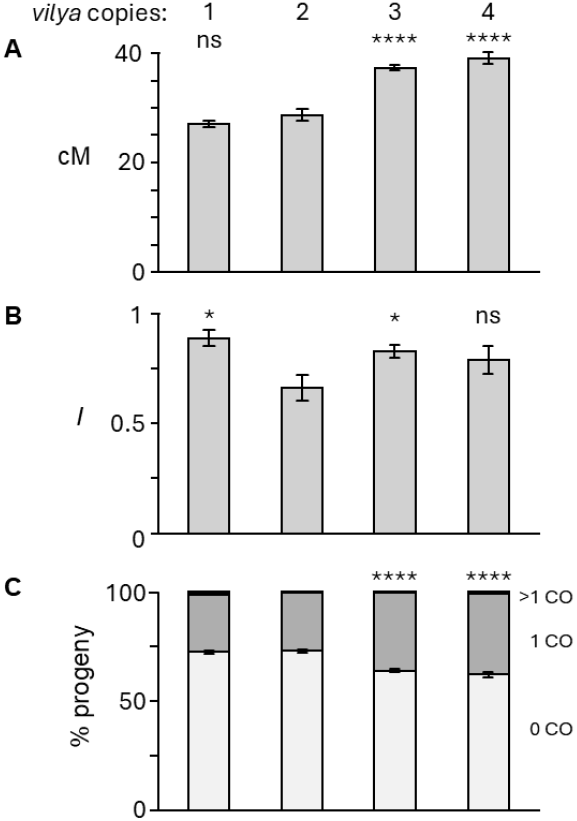
Figure 1 Effects of *vilya* dosage on recombination and interference. A) Crossovers in flies with different *vilya* dosage. Bars show the genetic length in centiMorgans (cM) of the region assayed in females with one, two (wild type), three, or four copies of *vilya*. Error bars are 95% confidence intervals. Statistical significance is indicated for comparisons to the wild type (two copies of *vilya*): ^****^: *p*<0.0001; ns: *p*>0.05. B) Interference values for different *vilya* dosages. Error bars are 95% confidence intervals. Statistical significance is indicated for comparisons to the wild type (two copies of *vilya*): ^*^: *p*<0.05; ns: *p*>0.05. C) Distributions of numbers of crossovers in progeny. Light gray, no crossovers; medium gray, one crossover; black, two or three crossovers. Error bars are 95% confidence intervals for the zero crossover category. Number of progeny scored (left to right) = 6805, 7666, 8692, and 1892.

We calculated Stevens’ measure of interference, *I*, where *I* = 1 indicates complete interference (no DCOs) and *I* = 0 means no interference (the number of DCOs observed is the same as the number expected if crossovers in each are independent of one another). We calculated *I* for the two larger, adjacent intervals. In the dataset from wild-type females, we expected 48 DCOs if there was no interference; we observed only 29 (*I* = 0.66±0.12; *p* < 0.0001). In flies heterozygous for a deletion of *vilya*, we expected 89 DCOs if there was no interference but observed only nine (*I* = 0.89±0.07; *p* < 0.0001). This is a significant reduction in interference in the *vilya* deletion heterozygote (*p* = 0.0282).

The significant decrease in DCOs in females heterozygous for a *vilya* deletion is consistent with Vilya playing a role in crossover designation through coarsening, because reduced Vilya would lead to a corresponding reduction in the number of condensates that lead to crossover designation. Since *2L* comprises about 20% of the euchromatic genome, reducing Vilya by 50% may leave a sufficient quantity to still have an average of one crossover designation per meiosis, while significantly decreasing the occurrence of two designations on *2L*. We note that among the 6805 progeny of wild-type females that we scored, 1801 had a single crossover in the interval assayed, and 64 had two crossovers. If each of the double-crossover meioses was changed to a single-crossover meiosis, the reduction in number of crossovers would not be statistically significant for this sample size.

### Crossovers are increased when there are additional copies of *vilya*

We also assayed the effects of increased Vilya dosage, using a duplication of *vilya* on an autosome. In females with one copy of the duplication, crossovers were significantly increased (37.1 cM in the region assayed, compared to 28.4 cM in wild-type females; *p* < 0.0001). There was no additional increase in crossovers in females homozygous for the duplication, and therefore carrying four copies of *vilya* (38.9 cM versus 37.1 cM, *p* = 0.15). It is possible that the system has become saturated for the amount of Vilya or that feedback on expression or protein degradation limits overexpression. If crossover designation by Vilya requires interaction with Narya and/or Nenya, then the amounts of Narya and Nenya may limit the effects of overexpression of Vilya alone.

In the coarsening model, overexpression of CORs would be expected to decrease crossover interference. This was not the case in our dataset from *vilya* duplication females. In females with a single copy of the duplication, interference was significantly increased (*I* = 0.834; *p* = 0.0155). Interference also appeared to be elevated in flies with two copies of the duplication (*I* = 0.788), but the difference was not statistically significant when compared to the other datasets (reduced fecundity of this genotype resulted in a smaller sample size). Increased crossing over may seem to be incongruent with interference being unchanged or increased. In wild-type females, 72.6% of the progeny had the parental configuration of markers and 26.5% had a chromosome with one crossover (Fig. 1C). In contrast, among the progeny of females with one copy of the duplication, 63.9% had no crossovers and 36.3% had a single crossover (*p* <0.0001). Thus, the elevated crossing over when Vilya dosage in increased is due to more chromosomes with one crossover and fewer with no crossovers (*p* <0.001). Because measures of interference require scoring large numbers of progeny, we assayed only about 50% of the genetic length of *2L*; measuring crossovers along an entire arm might lead to additional insights.

### Crossovers are increased when all three CORs are overexpressed

*C. elegans* ZHP-3 and ZHP-4 interact physically (Nguyen et al., 2018; Zhang et al., 2018). In a yeast two-hybrid assay, Vilya, Narya, and Nenya each interact with themselves and with each other (Lake et al., 2019). We therefore used the Gal4-*UAS* system to overexpress all three proteins coordinately. We constructed a fusion of a *10xUAS::hsp70z* promoter to the open reading frames encoding each COR, then assembled a plasmid carrying all three, a *y*^*+*^ gene as a marker for transformation, and a phiC31 *attB* site for integration into a genomic *attP* landing site. We scored crossovers in females carrying this transgene and either *bam::GAL4* or *nos::GAL4*, both of which express Gal4 in the female germline beginning before pachytene (Rørth, 1998). The genetic length of the test interval increased from 28.4 cM in wild-type females to 31.5 cM in those carrying the *bam::GAL4* transgene (*p* <0.0001) and to 32.1 cM in those with the *nos::GAL4* transgene (*p* <0.0001) (Figure 2A). There was no significant change in interference, although the only two triple crossovers we saw were both in the *bam::GAL4* progeny. As was the case for the *vilya* duplication, the increased crossovers in the *bam::GAL4* or *nos::GAL4* experiments are observed primarily as an increase in single-crossover chromosomes (Fig. 2C). We conclude that overexpression of the three COR proteins in meiosis results in a significant increase in crossovers.

**Figure 2.**
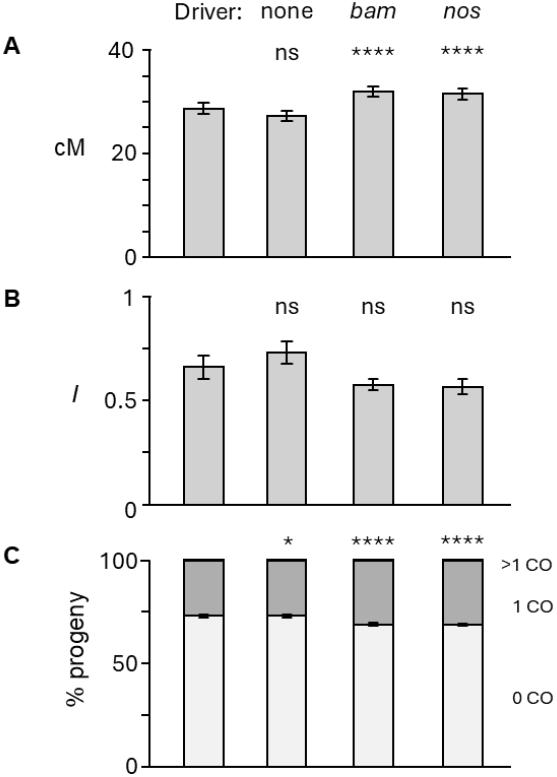
Figure 2 Crossing over when Vilya, Narya, and Nenya are coordinately overexpressed. Bars on the left are the data from wild-type females from Figure 1 (2nd from left in that figure). The second set of bars are from flies carrying the transgene with *UAS::vilya, UAS::narya*, and *UAS::nenya*, but no Gal4 driver, and those on the right are the same but with *bam::GAL4* or *nos::GAL4*. A) Crossovers in the assayed region. Error bars are 95% confidence intervals. For statistical comparisons in all panels, the no driver control was compared to wild-type females, whereas the *bam::GAL4* and *nos::GAL4* were compared to the no driver control. ns: *p*>0.05; ^****^: *p*<0.0001. B) Interference values for different *vilya* dosages. Error bars are 95% confidence intervals. C) Distributions of numbers of crossovers in progeny. Light gray, no crossovers; medium gray, one crossover; black, two or three crossovers. Error bars are 95% confidence intervals for the zero crossover category. Number of progeny scored (left to right) = 7666, 5251, 4911, and 7551.

### Functions of CORs in Drosophila recombination

Previous research using a combination of mutations and RNAi knockdown indicated that the Drosophila CORs are required both for DSB formation and for crossover generation (Lake et al., 2015, 2019). Narya and Neyna appear to be redundant with one another in these experiments. However, given that the gene duplication that generated *narya* occurred about 10-12 million years ago (Kumar et al., 2022), and both *narya* and *nenya* are apparently intact in all nine species in the melanogaster subgroup, it seems likely that each has some important unique function. Alternatively, expression of both might be required to achieve a sufficient concentration to ensure a least one crossover designation per major chromosome arm per meiosis.

Cytological studies show that Vilya is distributed through the SC but is enriched at sites of DSBs early in pachytene, then later is found at foci that are similar in number per nucleus to the number of crossovers, and are typically one per arm (Lake et al., 2015, 2019). These observations are consistent with the coarsening model. The findings we describe here on effects of altering COR dosage further support this model.

Crossover designation through coarsening may explain the longstanding puzzle of why there are no meiotic crossover on *Drosophila melanogaster* chromosome *4*. The finding that mutants that lack the anti-crossover protein Blm have crossovers on *4* (Hatkevich et al., 2017) suggested that the absence of crossovers in wildtype flies is due to active suppression by crossover patterning processes. The mechanism that suppresses crossovers in centromere-proximal regions seemed like a good candidate (Hartmann and Sekelsky, 2017); however, subsequent studies found that crossovers do occur in regions of similar size and distance from the centromere on *2R* (Hartmann et al., 2019; Pazhayam et al., 2024, 2025). We propose instead that the SC of *4* is too short to harbor enough CORs to achieve an accumulation that will lead to a crossover designation. The other five chromosome arms are each more than five times the length of *4* (including the pericentromeric heterochromatic satellite sequence, which may or may not participate in coarsening) and each averages just over one crossover per meiosis. It is unlikely that increased expression of COR proteins as we describe here would overcome this limitation.

## Methods

### Drosophila genetics

Flies were raised on standard medium purchased from Archon Scientific (Durham, NC). Deletion and duplication stocks were obtained from the Bloomington Drosophila Stock Center. The deletion used to reduce *vilya* dosage was *Df(1)ED6630* (RRID:BDSC_8948), which removes a total of 41 genes. The duplication used was *Dp(1;3)DC406* (RRID:BDSC_31456). These and other genetic elements are described in Flybase (release FB2025_05, (Öztürk-Çolak et al., 2024)).

### Transgene construction and transformation

The UAS.VNN transgene was constructed using GoldenBraid cloning (Sarrion-Perdigones et al., 2013; Matinyan et al., 2021). Vectors and some GoldenBraid parts were obtained from Addgene, and parts we made were deposited into Addgene; accession numbers are in parentheses. A transcription unit (TU) for each COR was assembled by combining a 10xUAS (252564), the hsp70z basal promoter (252565), the Vilya, Narya, or Nenya open reading frame (intronless coding sequences flanked by sequences for cloning into pUPD2 were synthesized as gBlocks by Integrated DNA Technologies, Inc.), and an HSV-tk terminator (165801). The Vilya and Narya TUs were assembled into Alpha1 (118044) to generate UAS.Vilya@A1 and UAS.Narya@A1. The Nenya TU was constructed in Alpha2 (118045) to generate UAS.Narya@A2. A *y*^*+*^ transgene was cloned into Alpha2 to generate y@A2 (252566). UAS.Vilya@A1 was combined with y@A2 into Omega1 (118046), and UAS.Narya@A2 was combined with a *bsr* gene in Alpha2 (derived from 165838) into Omega2 (118047). The resulting UAS.Vilya+y@O1 and UAS.Narya+bsr@O2 were combined back into Alpha1 to make UAS.Vilya+y+UAS.Narya+bsr@A1. This was combined with UAS.Nenya@A2 into Omega1.attB, which is an Omega1 vector with a *phiC31 attB* site inserted (252567). The resulting plasmid had UAS::Vilya, y+, UAS::Narya, bsr, UAS::Nenya, and the *attB* site. This plasmid was sent to GenetiVision (Houston, TX) for injection into VK31, a strain with a phiC31 *attP* insertion in 62E1. Adults that developed from injected embryos were crossed to *y w* ; *TM3, Sb*/*TM6B, Tb Hu*, and progeny were screened for expression of the *y*^*+*^ marker. Integrations were confirmed by PCR.

### Recombination and interference

To measure crossing over on *2L* we made females heterozygous for a chromosome carrying *dpp*^*d-ho*^ *dpy* (unknown allele) and *Adc*^*b-1*^ (referred to hereafter with the classical gene name *b*) and a chromosome carrying *P*{*w*^*+*^}*KG03050*, which is inserted about midway between *dpy* and *b*. Virgin females heterozygous for these two chromosomes were backcrossed to *w*^*1118*^; *dpp*^*d-ho*^ *dpy b*^*1*^ males and the relevant phenotypes (heldout wings versus normal posture wings; dumpy wings versus normal shape wings; red eyes versus white eyes; and black body versus normal body color) were scored in the progeny. Genetic distances are expressed here using the more common centiMorgan (cM) unit rather than the equivalent map units traditionally used in *Drosophila*. Measures of interference were calculated using the equations of Stevens (1936). The coefficient of coincidence (*c*) is calculated as 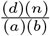, where *a* and *b* are the number of single-crossover progeny in the two intervals being compared, *d* is the number of double-crossover progeny, and *n* is the total number of progeny scored. This is equivalent to observed DCOs divided by expected DCOs if there is no interference. Interference (*I*) is equal to 1-*c*, and typically ranges from 0 (no interference) to 1 (complete positive interference, meaning no DCO progeny).

### Statistical analyses

For cM, *f* (crossover fraction) is the number of crossovers divided by total progeny scored (*n*). Variance (*V*) is *f* (1-*f*)/*n*, and standard deviation (SD) is 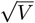. 95% confidence intervals were calculated as SD^*^1.96. For comparisons of crossover numbers in two genotypes, we used GraphPad QuickCalcs to conduct chi-square tests with Yate’s continuity correction. To compare interference values, we did Fisher’s exact tests using observed and expected number of DCOs for each genotype. For distributions of parental, single crossover, and double crossover classes, we did chisquare tests between observed and expected, where expected was the number expected in a given sample size if the distribution between classes was the same as in the control.

### Data availability

Raw counts of progeny classes per vial can be found at figshare.com under doi 10.6084/m9.figshare.31096369.

## Acknowledgments

We thank members of the Sekelsky lab for comments on the manuscript. Stocks obtained from the Bloomington Drosophila Stock Center (NIH P40OD018537) were used in this study.

## Funding

This work was supported by a grant from the National Institute of General Medical Sciences to JS under award 1R35GM118127 and by a Summer Undergraduate Research Fellowship to EF from the Office for Undergraduate Research at the University of North Carolina at Chapel Hill.

## Conflicts of interest

The authors declare that they have no conflicts of interest.

## Notes

https:figshare.com/10.6084/m9.figshare.31096369

### Competing Interest Statement

The authors have declared no competing interest.

